# Parvalbumin cell ribosome profiling in adult enhanced plasticity paradigms reveals distinct molecular signatures compared to juvenile critical period plasticity

**DOI:** 10.1101/2023.09.11.557035

**Authors:** David Benacom, Camille Chataing, Jérôme Ribot, Isabelle Queguiner, Alain Prochiantz, Ariel A. Di Nardo

## Abstract

The juvenile brain is shaped by critical periods (CPs) of plasticity regulated in part by parvalbumin (PV) interneurons. The “reopening” of CPs has been explored in brain disorders, yet it remains unclear whether adult enhanced plasticity recapitulates developmental mechanisms. We profiled visual cortex PV-specific ribosome-associated mRNAs at the CP peak and in two adult plasticity paradigms, chondroitinase ABC (ChABC) injection and extracellular OTX2 blockade. No shared translatomic signature emerged between the three conditions. Nonetheless, CP and ChABC-induced plasticity shared genes coding for synaptic components and extracellular matrix. Condition-specific pathways highlighted mechano-transduction and cytoskeletal remodeling for ChABC plasticity and DNA repair and heterochromatin for CP plasticity. These juvenile pathways were supported by elevated γH2AX, H3K9me3, and DAPI foci in PV cells uniquely during the CP. Thus, adult plasticity does not simply reinstate juvenile mechanisms, but instead may recruit partially overlapping, condition-specific molecular pathways revealing candidate targets for enhancing adult plasticity.

## Introduction

Neural circuits are strongly shaped by experience during restricted windows of postnatal plasticity known as critical periods (CPs), when sensory inputs can durably reconfigure connectivity^1–3^. Primary visual cortex (V1) is a canonical model for dissecting the underlying mechanisms: monocular deprivation (MD) experiments revealed that ocular dominance (OD) rewiring is extensive and rapid only during a juvenile CP and is strongly attenuated in adulthood, providing an experimental paradigm to relate cellular, synaptic, and molecular events to network plasticity.

Fast-spiking parvalbumin (PV) interneurons in layers III–IV of the cerebral cortices have emerged as central CP regulators^4^. Their molecular maturation and the associated increase in principal cell perisomatic inhibition are tightly coupled to CP onset. These molecular changes are widespread, ranging from the epigenetic level, with changes in histone acetylation and DNA methylation, to the extracellular level, with changes in extracellular matrix (ECM) composition and the accumulation of dense perineuronal nets (PNNs) around PV cell soma and proximal dendrites. PNNs provide structural restriction of plasticity^5–7^ and are involved in signaling through the specific recognition of molecular cues^8^, such as the homeoprotein transcription factor OTX2 released from the choroid plexus into cerebrospinal fluid^9^. Following its binding to PNNs, OTX2 is internalized by PV cells and participates in regulating both CP opening and closure^10^. CPs also involve the contribution of glial cells^11^: astrocytes regulate ECM composition and PNN integrity^12^, oligodendrocytes myelinate PV cells^13^, and microglia prune and regulate synaptic structures^14^. At CP closure, these converging processes establish multiple, partially redundant, molecular “brakes” on circuit remodeling, rendering adult V1 relatively resistant to perturbations in sensory input.

Nevertheless, the adult cortex retains a latent capacity for plasticity^15^. Local treatment with chondroitinase ABC (ChABC) in adult V1 enzymatically disrupts PNNs and restores OD plasticity^16^. Conditional expression of a secreted anti-OTX2 single-chain antibody (scFv-OTX2) in cerebrospinal fluid neutralizes extracellular OTX2 and antagonizes its uptake by PV cells, resulting in reduced PV and PNN levels in adult V1 and in functional OD plasticity^17^. Finally, the treatment with HDAC inhibitors reactivates adult OD plasticity by increasing accessibility to enhancer regions^18^. The implication of PNNs, histone acetylation and OTX2 in both CP mechanisms and adult plasticity paradigms suggests that there are fundamental molecular similarities between CPs and therapeutic plasticity.

To further address to what degree enhanced plasticity in the adult resembles juvenile CP plasticity, we chose to focus on PV cells given their central role in CP regulation. PV-cell-specific ribosome-bound mRNAs (hereafter referred to as translatome) were isolated by translating ribosome affinity purification and quantified by RNA sequencing (TRAP-seq). This approach captures mRNAs that are engaged in active translation, providing a proximal readout of the proteins being synthesized throughout the cell, from soma to synapse, reflecting the molecular dynamics driving cellular activity. PV cell translatomes were profiled in V1 under five conditions: peak endogenous CP in juvenile mice; adult “non-plastic” state; adult plasticity after ChABC treatment; adult plasticity after local expression of secreted scFv-OTX2; and aged (>2 yr-old) adult mice for which there is a potential for reduced PNN levels and dysregulated circuits in sensory cortices^19,20^.

## Results

### PV and PNN levels are consistent across CP and adult V1 plasticity paradigms

We first evaluated whether the levels of general PV cell markers change as expected in juvenile V1 during CP plasticity and in adult V1 after induction of enhanced plasticity. Coronal sections were stained for PV and for PNNs by using *Wisteria floribunda* lectin (WFA). Postnatal day (P) 90 was used a “non-plastic” adult age for reference to compare with peak CP at P30 and with two enhanced-plasticity paradigms at P90 involving either ChABC injection or scFv-OTX2 expression in V1. A pre-trained, fine-tuned deep-learning model was used for unbiased quantification of cell number and staining intensity (see Methods). Wild-type mice showed a small increase in PV^+^ cell number along with large increases in PV staining intensity and double-stained WFA^+^PV^+^ cell number at P90 compared to P30 (Fig. 1a). To enhance plasticity via PNN removal, ChABC was injected unilaterally in adult V1, and PV markers were evaluated 4 d later, by which time PNN levels are known to be decreased and have yet to recover (Fig. 1b)^21,22^. A steep loss of WFA^+^ cells confirmed PNN disruption, which was accompanied by a decrease in the number of PV^+^ cells, with no change in PV staining intensity, as reported previously^23^. For the scFv-OTX2 condition, restricted expression in PV cells was obtained by unilateral injection of an adeno-associated virus (AAV) encoding Cre-dependent scFv-OTX2 in V1 of P70 mice. Extracellular OTX2 neutralization led to significant PNN loss in the injected hemisphere at P90 and was accompanied by a non-significant decrease in PV cell number and staining intensity (Fig. 1c).

**Fig. 1.**
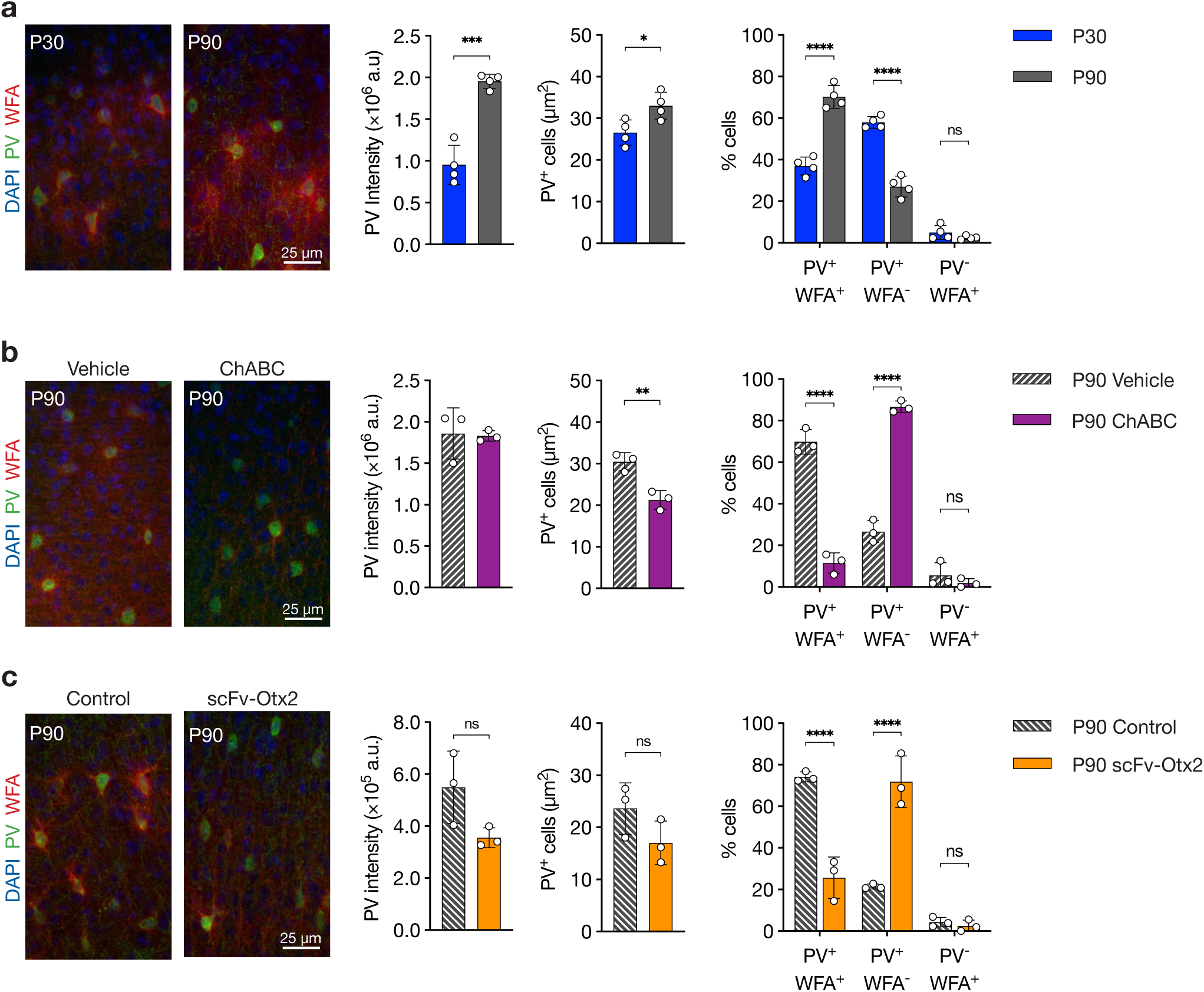
PV cell morphology is consistent with V1 plasticity state. **a**, Representative images of V1 supragranular layers at P30 and P90 stained for PV and WFA, with quantification of PV intensity, PV^+^ cell density, and co-localization of PV^+^ and WFA^+^ cells. Unpaired *t*-test; n per group = 4. **b,** Representative images of V1 supragranular layers at P90 of vehicle- and ChABC-injected cortex stained for PV and WFA, with quantification of PV intensity, PV^+^ cell density, and co-localization of PV^+^ and WFA^+^ cells. Unpaired *t*-test; n per group = 3. **c,** Representative images of V1 supragranular layers at P90 of control and scFv-Otx2-expressing cortex stained for PV and WFA, with quantification of PV intensity, PV^+^ cell density, and WFA^+^ cell density. Paired *t*-test; n per group = 3. All values ± SD; * *P* < 0.05, ** *P*<0.01, *** *P* < 0.001, **** *P* < 0.0001; ns, non-significant.

Since adult enhanced plasticity induced by extracellular OTX2 neutralization was only previously demonstrated following scFv-OTX2 secretion from the choroid plexus^17^, we verified whether V1 plasticity occurs after its local expression and secretion by V1 PV cells. To that end, 3 weeks after AAV injection, mice were submitted to 3-day MD followed by optical imaging of blood-oxygen-level-dependent (BOLD) response in ipsilateral and contralateral V1 (Fig. 2a,b). Functional plasticity was observed with an OD index shift towards ipsilateral V1 (Fig. 2c) driven by increased ipsilateral activation (Fig. 2d). To determine the effect of local scFv-OTX2 expression on visual acuity, the visual water task and the visual cliff test were used (Fig. 2e). Both assessments showed no difference between control and scFv-OTX2-treated mice, confirming that visual acuity remained unaltered (Fig. 2f,g). The observed OD shift is thus not linked to a loss of visual acuity following local extracellular OTX2 neutralization.

**Fig. 2.**
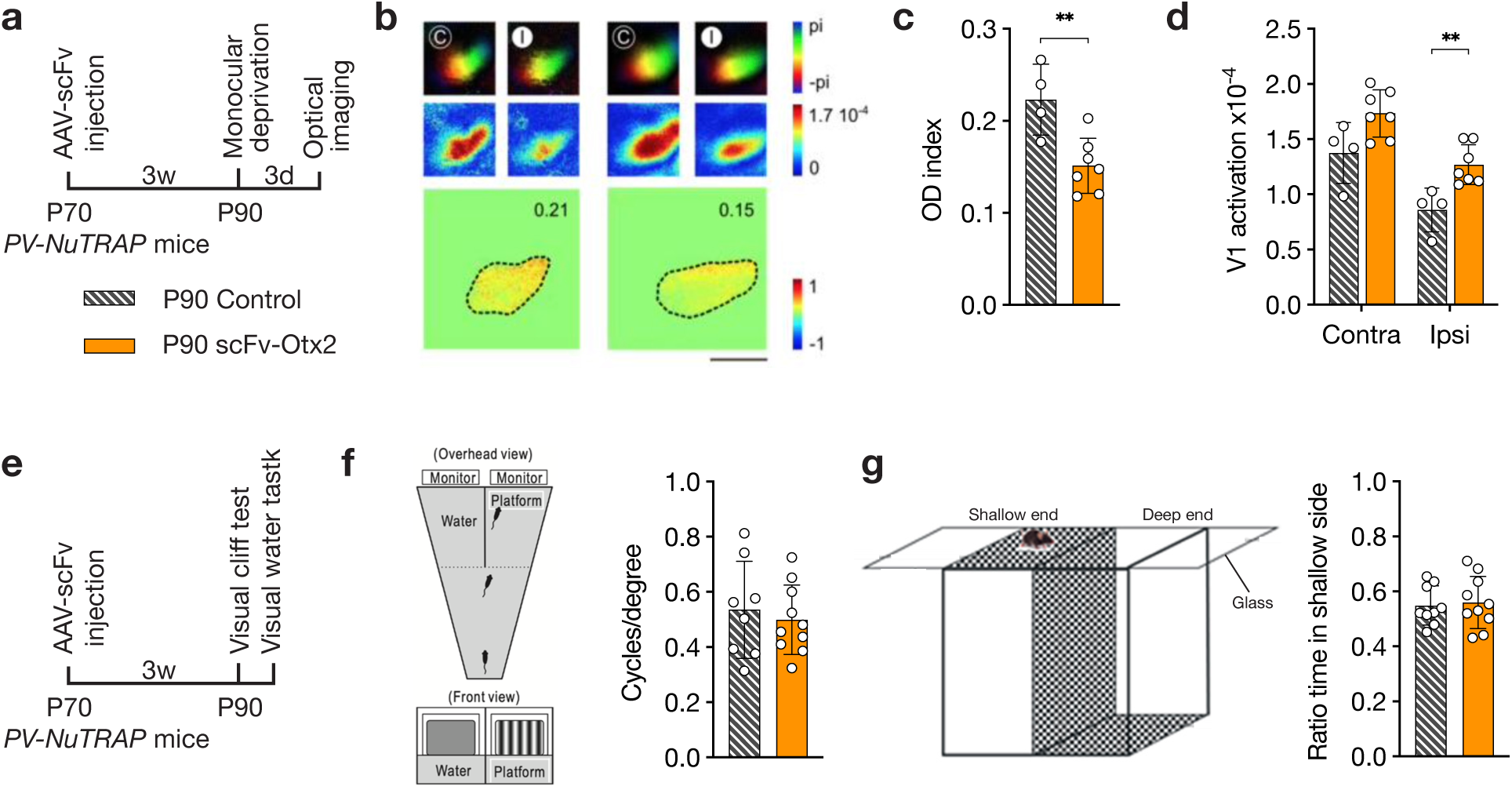
Adult expression of scFv-Otx2 secreted from PV cells leads to functional plasticity with no change in visual acuity. **a,** Timeline for stereotaxic surgery and ocular dominance (OD) analysis of *PV-NuTRAP* mice expressing secreted scFv-Otx2 in V1 PV cells. **b,** Visual maps of control mice and mice expressing scFv-Otx2. Contralateral (C) and ipsilateral (I) visual stimulations are indicated with a white (non-deprived eye) or black (deprived eye) circle. The bottom panels show the normalized OD map with the average value inset. **c,** OD index calculated from visual maps. Unpaired *t*-test; n per group = 4 or 7. **d,** Relative activation of V1 in contralateral (Contra) and ipsilateral (Ipsi) hemispheres. 2-way ANOVA; n per group = 4 or 7. **e,** Timeline for stereotaxic surgery and acuity analysis of PV-NuTRAP mice expressing secreted scFv-Otx2 in V1 PV cells. **f,** Visual water task requires mice to swim to side with randomly-placed platform indicated by a monitor presenting gratings with variable spatial frequency. Visual acuity of each mouse was determined by averaging the maximal performance of the last 20 trials. Unpaired *t*-test; n per group = 9 or 10. **g,** The visual cliff test to determine depth perception was quantified by the ratio of time spent in the shallow side versus the deep side. Unpaired *t*-test; n per group = 9 or 10. All values ± SD; ** *P*<0.01.

### V1 PV cell translatomes reveal the absence of a plasticity-specific signature across all experimental paradigms

To evaluate whether PV cell translation in mouse V1 is similar during juvenile OD CP plasticity and after plasticity has been enhanced in the adult, *PV-NuTRAP* mice were used for immunoprecipitation (IP) of GFP-labeled ribosomes to isolate actively translated ribosome-bound RNA, hereafter termed PV-IP fraction. Double staining for PV and GFP on V1 sections at P30 showed that nearly all PV^+^ cells co-express GFP, while around 80% of GFP^+^ cells co-express PV (Fig. 3a). The absence of PV staining in certain GFP^+^ cells is likely due to low-level PV expression as previously reported^24^. A TRAP experiment was performed on V1 lysates at P30, P90, and in adult ChABC and scFv-OTX2 experimental paradigms. Aged mice (P800) were also included to compare features of juvenile and adult-enhanced plasticity with putative age-associated changes. For each condition, RNA from V1 lysates (input) and TRAP samples (PV-IP) were sequenced and for each technical replicate, lysates from 3 to 5 mice were combined in each sample to obtain enough material and to mitigate biological variation (Fig. 3b). Principal component (PC) analysis revealed high-quality clustering of samples in both input and PV-IP fractions (Fig. 3c,d). The high quality of the pulldown was confirmed by comparing RNA levels between input and PV-IP fractions of markers for other neuronal types and glial cells (Fig. 3e). *Pvalb* was the most enriched mRNA, and several other mRNA were identified as PV-specific (Fig. 3e,f). Genes highly downregulated in PV-IP, comprising glial cell markers were removed from the PV-IP analysis (see Methods). Comparison of PV-IP to input fractions across all experimental conditions identified ∼1600 genes systematically enriched (log_2_FC > 0.3) in the PV-IP fraction of each condition (i.e., the intersection of all sets), as well as ∼4500 genes that were enriched in at least one experimental condition (i.e., the union of all sets) (Fig. 3g). We considered the union of all PV-IP sets as putative PV-enriched transcripts for downstream analysis.

**Fig. 3.**
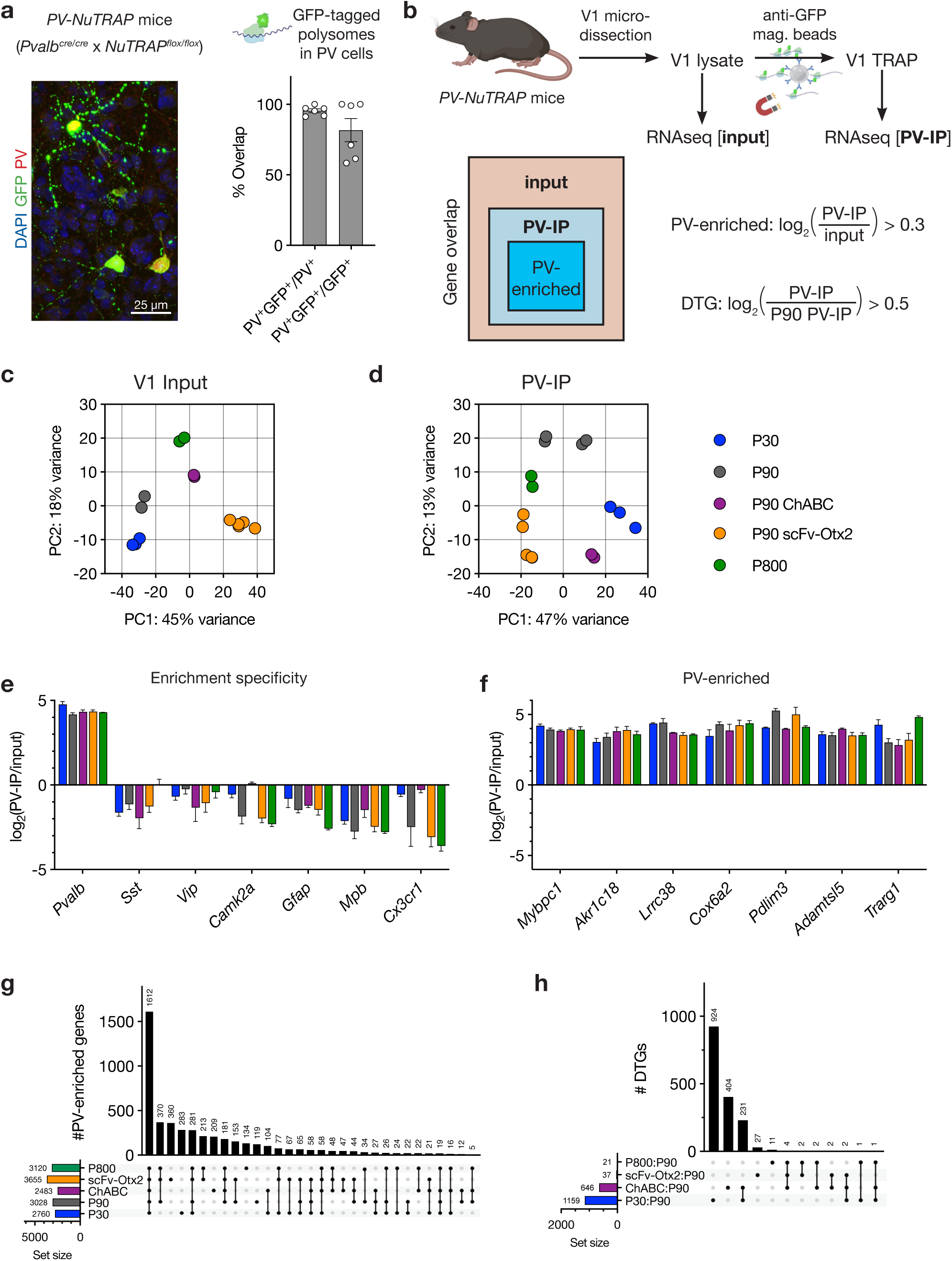
TRAP analysis of V1 PV cells reveals plasticity-dependent changes. **a,** Validation of the PV-NuTRAP mouse model for constitutive expression of ribosome-associated GFP in PV cells. Representative image of supragranular cortex stained for PV and GFP along with quantification of double-stained cells. **b,** RNA-seq was performed on *PV-NuTRAP* mouse V1 lysates (input) and V1 TRAP (PV-IP) samples. Comparison of PV-IP values to input values provided a list of PV-enriched genes, while differentially translated genes (DTGs) were identified by comparing PV-IP values to P90 PV-IP values. **c,** Principal component (PC) analysis of V1 input samples. **d,** Principal component analysis of PV-IP samples. **e,** PV-enrichment values for genes representative of cortical excitatory and inhibitory neurons and glial cells. **f,** Enrichment values of the most highly PV-enriched genes across all groups. **g-h,** Upset plots of the number of PV-IP genes enriched in PV cells (g) and PV-enriched DTGs (h) showing overlap in enriched genes between the various groups. Black dots indicate the presence of genes within a given group.

To determine whether a common translation signature underlying plastic-like states exists, differentially translated genes (DTGs) in the PV-IP fractions were calculated by comparing to the control P90 PV-IP fraction (Fig. 3h). The comparison of P30:P90 yielded the highest number of DTGs, with 1159 in total (831 upregulated and 328 downregulated), of which 497 were PV-enriched (270 up and 227 down). The ChABC:P90 comparison showed 646 DTGs overall (487 up and 159 down), including 303 PV-enriched genes (189 up and 114 down). The scFv-OTX2:P90 comparison also showed very limited changes, with only 37 DTGs (33 up and 4 down) and only 6 PV-enriched transcripts (4 up and 2 down), suggesting that scFv-OTX2-induced plasticity relies on minimal translational remodeling within PV cells. The aged condition P800:P90 showed a smaller but detectable modulation, with 21 DTGs in total (13 up and 8 down), including 10 PV-enriched (5 up and 5 down), suggesting a very limited reorganization of the PV translatome during aging. Strikingly, only two DTGs (*Vps13a* and *Cd63*) were found common to all three plastic-like conditions (P30, P90 ChABC, and P90 scFv-OTX2), suggesting that a common PV translatome signature is not associated with these three functional plasticity paradigms.

To represent the contribution of other V1 cells, we compared the fold-change of differentially expressed genes (DEGs) of the bulk input fractions to the fold-change of DTGs of PV-IP fractions using P90 as a reference (Fig. 4). As shown in other TRAP studies, the resulting diagonal plots helps determine if fold-changes are driven by the input fraction or by the cell-type of interest^25^. Diagonal plots reveal that P30 (Fig. 4a) and P90 ChABC (Fig. 4b) values are driven by changes in PV-IP fractions, that P90 scFv-OTX2 (Fig. 4c) is driven by the input fraction, and that P800 (Fig. 4d) has a similar number of DTGs compared to DEGs, suggesting changes are balanced across all cortical cells. Given the more widespread changes in the PV-IP fractions of both P30 and P90 ChABC, we focused on these conditions to further compare juvenile CP plasticity and adult enhanced plasticity.

**Fig. 4.**
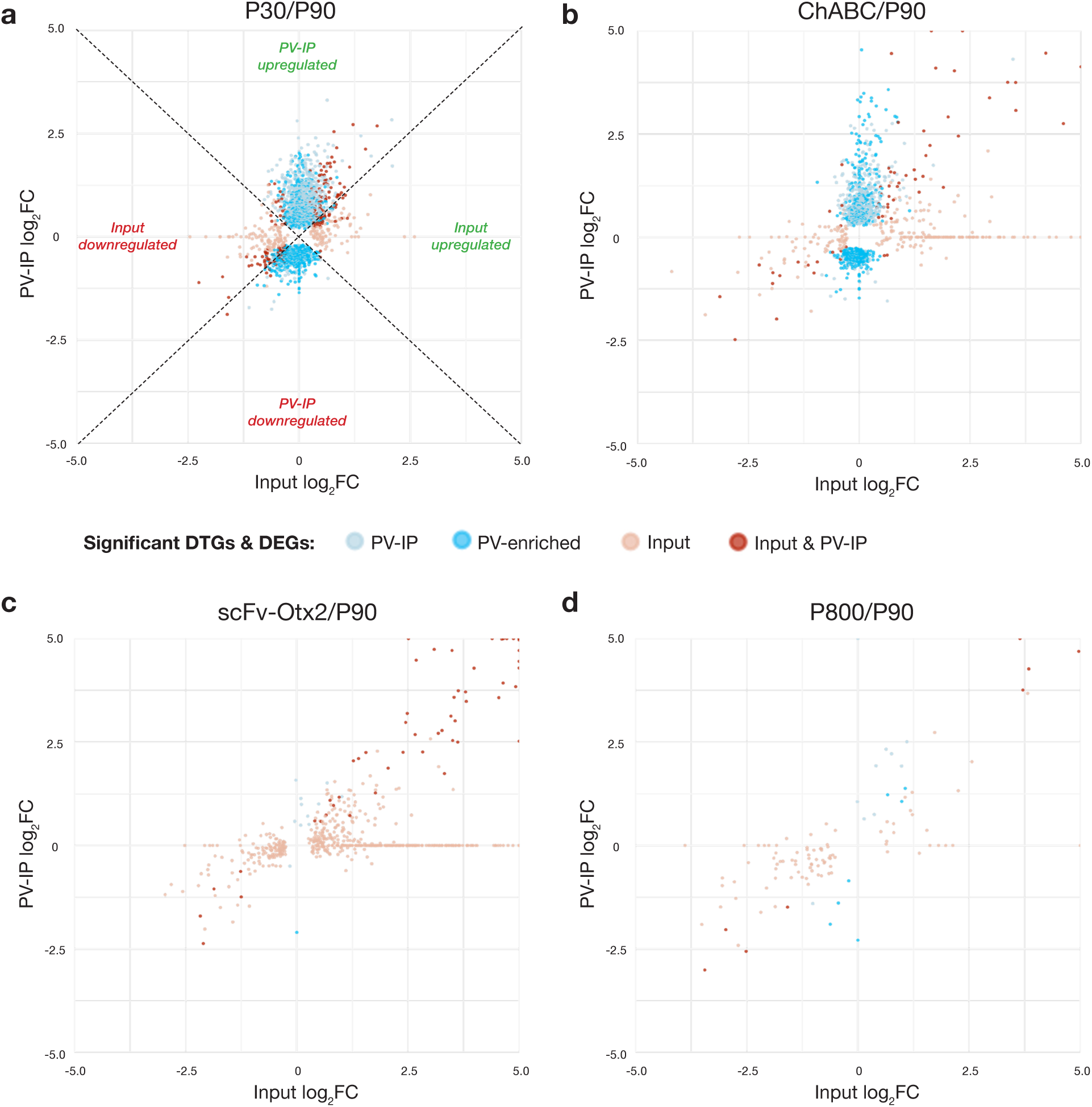
Translation is more impacted in PV cells at P30 and P90 ChABC compared to P90 scFv-Otx2 and aged mice. **a-d,** Diagonal plots comparing V1 PV cell DTGs (from PV-IP samples) and total DEGs (from input samples) for P30:P90 (a), ChABC:P90 (b), scFv-Otx2:P90 (c), and P800:P90 (d) groups. Only genes with FDR < 0.05 are shown. Genes along the diagonal are affected in both samples, genes along the horizontal are affected in the input samples, while genes along the vertical are affected in the PV-IP samples.

### V1 PV cell translatomes show common and condition-specific pathways between juvenile CP and adult ChABC-induced plasticity

To determine the level of similarity between physiological juvenile CP plasticity and adult ChABC-induced plasticity in PV cells, DTGs from P30 and P90 ChABC (n=1155 genes and n = 635 genes, respectively, excluding the ones in common with scFV or P800) were compared in depth (Fig. 5a). The intersection (P30:P90 ∩ ChABC:P90) identified a shared set of 234 genes (of which 90 were PV-enriched) that we defined as ‘common’ genes. A vast majority of these common DTGs displayed identical up or down directions relative to P90, whereas only a small subset (9 of 234) displayed opposing directions (i.e., up in ChABC:P90 and down in P30:P90). A hierarchical topGO analysis was performed separately on specific and common DTGs to identify cellular component (CC), molecular function (MF) and biological pathways (BP). The common DTGs highlighted synaptic plasticity and brain development (Fig. 5a), supported by pathways such as synaptic vesicle (CC), receptors (CC), synaptic receptors (MF), calmodulin binding (MF), extracellular space (CC), and establishment or maintenance of epithelial cell apical/basal polarity (BP) among other terms. A similar increase in several DTGs related to calcium-dependent activity and extracellular activity, for example (Supplementary Fig. 1b), emphasizes certain commonalities between P30 and P90 ChABC. Together, these pathways point towards a core plasticity program in both conditions that coordinates synaptic organization, receptor trafficking, ion channel regulation, and extracellular matrix remodeling. Interestingly, *Mef2c*, a master regulator of PV cell development^26,27^, was upregulated in P30 and P90 ChABC, suggesting a common pathway with developmental underpinnings (Supplementary Fig. 1c).

**Fig. 5.**
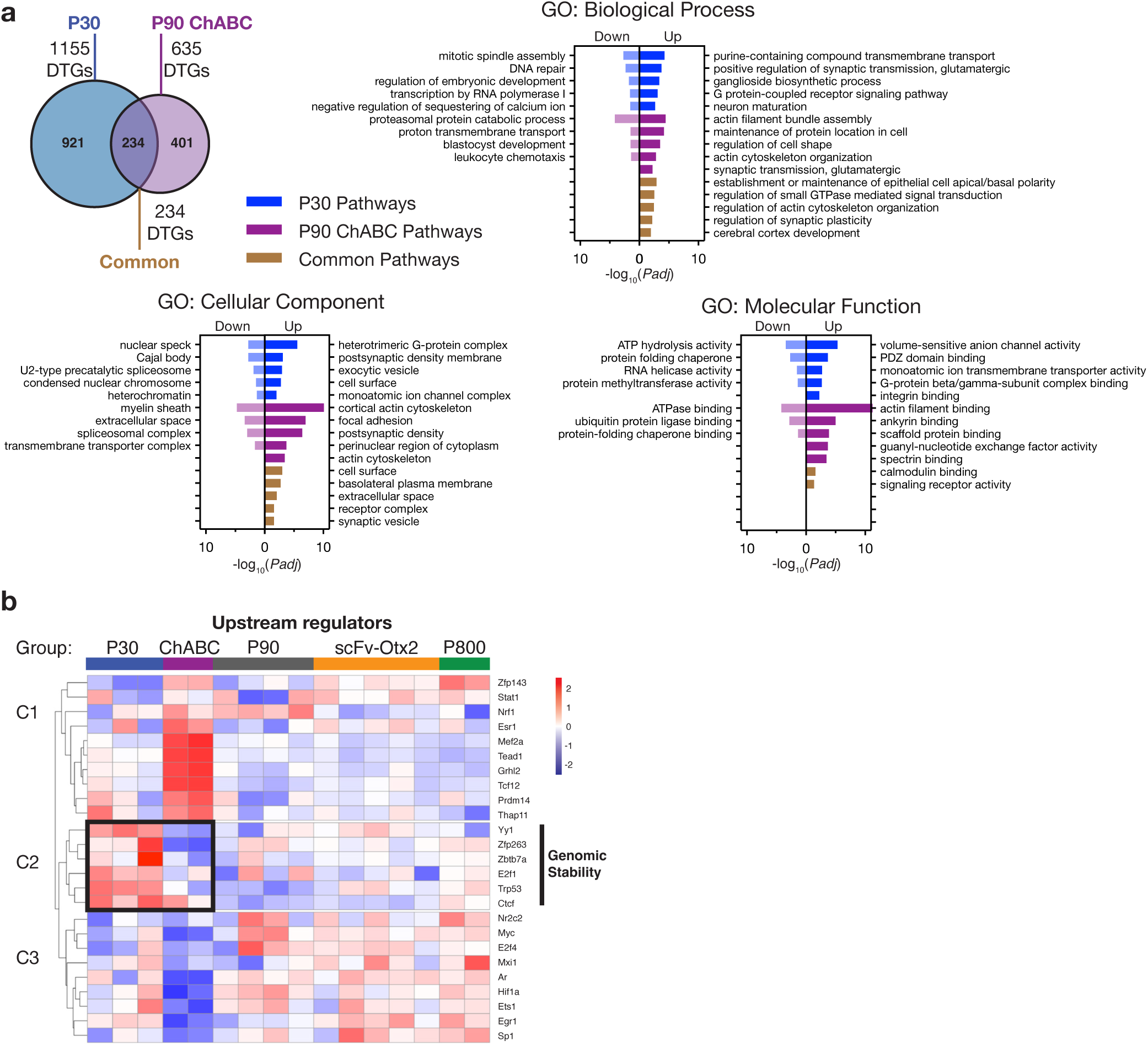
Pathway and transcription factor activity analysis reveal common and distinct outcomes between juvenile CP plasticity and adult enhanced plasticity. **a,** GO pathway analysis of DTGs common to P30 and P90 ChABC-treated V1, and DTGs specific for each condition. **b,** Cluster map of global transcription factor (TF) activity patterns based on PV-IP levels in each group: P30, P90, P90 ChABC, P90 scFv-Otx2, and P800. Black rectangle highlights opposing patterns in P30 and P90 ChABC in cluster C2 containing TFs related to genomic stability.

While juvenile CP and ChABC-induced adult plasticity share several pathways, we find clear evidence of other non-overlapping molecular responses as evidenced by condition-specific pathways (i.e., regulated in ChABC:P90 but not in P30:P90 and vice-versa). Analysis of the 401 ChABC-specific DTGs (of which 210 were PV-enriched) revealed pathways involved in cell shape (BP) and cytoskeleton regulation related to synaptic scaffolding (Fig. 5a). The expression profiles of the genes belonging to the scaffolding (MF: *Shank1, Shank3*…), ankyrin (MF: *Ank2, Ank3*…) and spectrin pathways (MF: *Sptb, Plec*…) confirm that these genes are not differentially translated in other conditions (Supplementary Fig. 1d,e). Regarding IEGs, *Fos* and *Fosb* were downregulated after ChABC treatment, consistent with previous findings (Supplementary Fig. 1c)^23^. Analysis of the 921 P30-specific DTGs (of which 407 were PV-enriched) revealed several upregulated pathways related to signaling and activity, such as voltage-gated potassium channel activity (MF; *Kcnn3, Kcnv1*…) and heterotrimeric G receptor (CC; *Gng12, Gnb4*…) (Fig. 5a and Supplementary Fig. 1f,g). Downregulated pathways point to broad nuclear changes at P30, involving DNA repair (BP), heterochromatin/nuclear speckles (CC) and methylase enzymes (MF)

### Upstream regulator analysis reveals condition-specific plasticity modules

To interpret condition-specific DTG patterns, upstream regulator analysis was performed on the PV-IP enriched fractions. Inferred transcription factor (TF) activities were found to be grouped into 3 clusters (Fig. 5b). Clusters C1 and C3 define the ChABC group with strong TF specificity. Cluster C1 points to upregulated activity of *Prdm14, Thap11, Esr1, Grhl2, Mef2a, Tead1, Tcf12,* and *Zfp143*, while C3 indicates downregulated activity of *E2f4*, *Myc*, *Hif1a*, *Ar*, and *Ets1*. The cluster C1 TFs are consistent with mechano-transduction, notably implicating TEAD/YAP–TAZ signaling, which translates ECM- and adhesion-dependent changes in actomyosin tension into mechanoresponsive transcriptional programs^28,29^. Cluster C2 contains P30-specific upregulated activity of *Zbtb7a, Yy1, E2f1, Ctcf* and *Trp53*, linking the observed DNA repair and heterochromatin pathways to known regulators of chromatin structure and genomic stability^30^. The identification of downregulated DNA-repair pathways along with a p53-centered TF cluster at P30 suggests a link between CP plasticity and DNA repair that is not recapitulated in adult ChABC-induced plasticity.

### Downregulated DNA repair and heterochromatin pathways at P30 are associated with increased γH2AX and H3K9me3 foci number in V1 PV cells

Compared to P90, only juvenile CP plasticity was associated with a downregulated DNA-repair pathway. Indeed, DTGs belonging to this pathway were found to be mostly unaltered at P90 ChABC (Fig. 6a). Several key components of canonical DNA double-strand break (DSB) response and DNA repair were markedly downregulated at P30, including members of the MRN complex (*Rad50, Mre11a*) involved in DSB sensing, the γH2AX-binding mediator *Mdc1*, and *Dmap1*, a regulator of ATM signaling. BER-associated factors (*Fen1, Xrcc1, Lig1*) were also downregulated at P30, together with chaperones supporting repair complex stability (*Hspa1a, Hspa1b*). To assess DNA-damage-associated chromatin changes in PV cells, γH2Ax and DAPI staining was performed in V1 at P30, P90, and at P90 with ChABC treatment (Fig. 6b). Automated quantification with a fine-tuned deep-learning model revealed a ∼2-fold higher number of γH2Ax foci in PV cells at P30 compared to P90, and P90 ChABC (Fig. 6c). Foci intensity and mean total nuclear intensity (including diffuse nuclear staining) showed that the numerous P30 foci were of lower γH2Ax intensity compared to the more condensed, larger foci appearing at P90 (Fig. 6d,e). Interestingly, the mean total nuclear γH2Ax intensity is also more elevated at P90 (Fig. 6e), suggesting that P30 and P90 PV cells have different chromatin contexts and/or DNA repair states, although their functional significance cannot be resolved from immunostaining alone. The DAPI heterochromatin-rich foci were also quantified in PV cells, showing the same pattern of a higher number of slightly less intense foci at P30 compared to P90 and P90 ChABC (Fig. 6c,d). However, mean total DAPI staining intensity remained constant between P30 and P90, which is coherent with DNA staining in post-mitotic neurons (Fig. 6e). Overall, these data suggest that the DNA repair pathways identified for juvenile CP plasticity are not recapitulated in ChABC-induced plasticity.

**Fig. 6.**
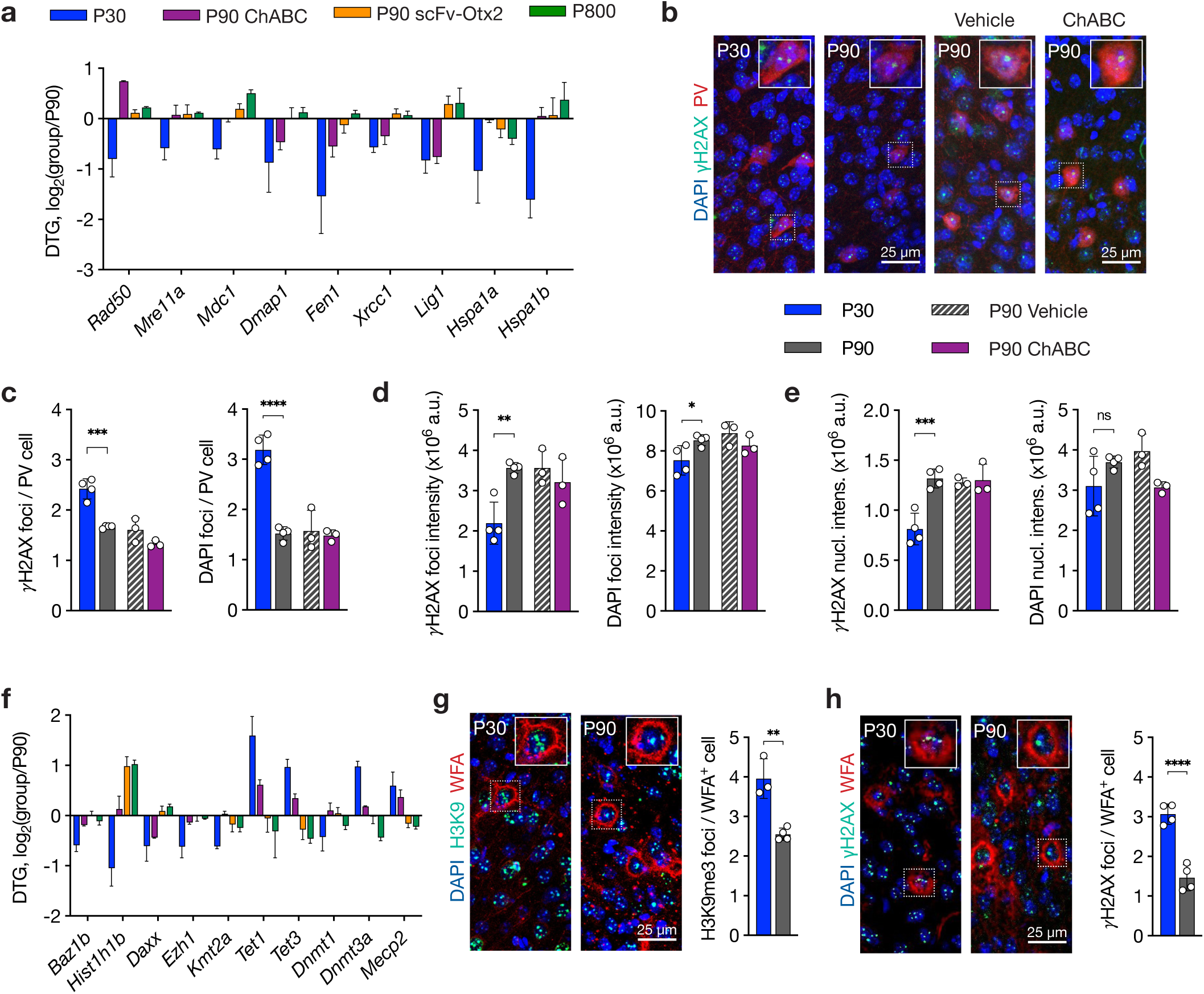
Decreased ribosome-association of DNA repair genes in V1 at P30 is accompanied by increased heterochromatin marks. **a,** Sample of DTGs involved in DNA repair showing decreases at P30 and minimal change in the other groups. **b,** Representative images of γH2AX and DAPI foci in PV cells in V1 of P30 and P90 mice, and P90 mice treated with ChABC or vehicle. **c-e,** Quantification of γH2AX and DAPI foci number (c), foci intensity (d), and mean nuclear staining intensity (e) in PV cells in V1 of P30 and P90 mice, and P90 mice treated with ChABC or vehicle. Unpaired *t*-test; n per group = 3 to 4. **f,** Sample of DTGs involved in heterochromatin structure and epigenetics. **g,** Representative images and quantification of H3K9me3 foci in WFA^+^ cells in V1 of P30 and P90 mice. Unpaired *t*-test; n per group = 3 to 4. **h,** Representative images and quantification of γH2AX foci in WFA^+^ cells in V1 of P30 and P90 mice. Unpaired *t*-test; n per group = 4. All values ± SD; * *P* < 0.05, ** *P*<0.01, *** *P* < 0.001, **** *P* < 0.0001.

The decreased number of large DAPI foci between P30 and P90 PV cells suggests general heterochromatin rearrangements, which is in keeping with the heterochromatin pathway identified in the P30 translatome. Indeed, DTGs at P30 included genes involved in chromatin organization (*Baz1b, Hist1h1b*), histone methylation (*Kmt2a, Ezh1*), de novo H3K9me3 deposition (*Daxx*), DNA methylation/demethylation machinery (*Tet1, Tet3, Dnmt1*, *Dnmt3a*) and the methyl-CpG-binding protein *Mecp2* (Fig. 6f). We previously reported that WFA^+^ cells, which represent the more mature PV cell population disrupted after ChABC treatment, have more MeCP2 foci at the peak of juvenile CP in V1 compared to adult^31^. Since MeCP2 binds to methylated DNA and can potentially direct the trimethylation of lysine 9 on histone H3 (H3K9me3), we further analyzed H3K9me3 foci at P30 and P90 in WFA^+^ cells. Their foci number was higher at P30, supporting developmental reorganization of H3K9me3-marked heterochromatin (Fig. 6g). While PV^+^ and WFA^+^ cells largely overlap (Fig. 1a), they might represent slightly different populations or PV cell maturation levels. Therefore, we also confirmed that γH2Ax foci number in WFA^+^ is also higher at P30 (Fig. 6h). Overall, these data support the DTGs and pathway-level changes identified by TRAP-seq, linking juvenile plasticity to altered DNA-damage-associated chromatin signaling and heterochromatin organization in PV cells.

## Discussion

Ribosome-associated mRNAs from PV cells in V1 were compared at P30, when juvenile OD CP plasticity is at its peak, at P90, long after CP closure, and at P90 in two well-established adult plasticity paradigms targeting PV cells: PNN hydrolysis by ChABC treatment and blockade of OTX2 uptake through a local expression of scFv-OTX2. While the latter paradigm enhanced functional OD plasticity in the adult mouse, it elicited very few DTGs in PV cells. This limited change in translated mRNAs was also observed in aged mice. In contrast, the P30 and P90 ChABC groups were associated with numerous and partly overlapping DTGs in PV cells. However, certain changes at P30, such as the downregulation of DNA repair and the re-arrangement of epigenetic marks, are not recapitulated in ChABC-induced adult plasticity.

The present study only focuses on ribosome-associated RNAs committed to translation throughout all cell compartments, including sites of local translation. Several prior studies have characterized molecular changes taking place during CP via bulk or single-nuclei (sn) RNA-seq^32,33^. Strikingly, snRNA-seq has revealed only mild PV cell changes before and after auditory CP in contrast to the dramatic physiological changes^33^. TRAP is not confined to the nuclear compartment and may therefore be more suitable for identifying transcripts reflecting mechanisms underlying plasticity^34,35^. These approaches are nonetheless complementary given that TRAP cannot survey regulatory layers involving miRNA and lncRNAs, for example.

A strong finding of this TRAP study is the absence of a common PV ribosome-associated signature between the three V1 plasticity paradigms. This illustrates that, compared to P90, functional plasticity can be obtained through distinct pathways that don’t necessarily “reopen” the CP in PV cells. The main divergence was found between P30 and P90 scFv-Otx2 plasticity, which share very few DTGs. In contrast, P30 and P90 ChABC display a certain overlap with concordant directions of change. These genes participate in classical plasticity pathways: synaptic organization, vesicle trafficking, cell adhesion, ECM and structural remodeling, and plasticity-dependent transcription regulation. However, some DTGs had opposing directions when comparing P30 and P90 ChABC, as is the case for *Rad50* mRNA. In other cases, DTGs specific to either P30 or P90 ChABC mapped to the same pathway, indicating that phenotypic convergence can be reached through different genetic mechanisms.

Beyond this commonality, several pathways are specific to the ChABC paradigm, with a clear emphasis on the cytoskeleton, including actin filament organization and ankyrin/spectrin-associated synaptic scaffolding modules. Upstream TFs indicate TEAD/YAP–TAZ signaling and mechanoresponsive transcriptional programs, which supports the idea that cytoskeletal and synaptic remodeling takes place downstream of ChABC-induced transcriptional response^36–38^. In addition, proteasomal and spliceosome pathways were identified in the ChABC-specific DTGs, suggesting that the relationship between ECM and neuronal programs extend beyond the disruption of surface receptors or synapses and provide an original source of future investigations in PNN-related biology.

Regarding the scFv-Otx2 plasticity paradigm, OTX2 import by PV cells is involved in both CP onset and closure according to a two-threshold model of OTX2 concentration^10^. Neutralizing extracellular OTX2 through scFv-OTX2 expression drops OTX2 levels below a stable threshold and “reopens” plasticity in the adult, raising the possibility that juvenile mechanisms are recapitulated. While, the present study confirms that local neutralization of extracellular OTX2 around PV cells provides functional plasticity in the adult, DTG analysis clearly demonstrates that the involved mechanisms differ from the juvenile CP. The paucity of DTGs in PV cells suggests that extracellular OTX2 neutralization engages downstream involvement of other cell types, including principal cells and astrocytes^12^, and/or engages different RNA species not captured by TRAP^39^.

At P30, V1 plasticity engages changes in PV cell epigenetic marks and in surface receptors likely representing the transition to fast-spiking activity. Voltage-gated potassium channels and G proteins complexes are specifically upregulated, suggesting that the channel repertoire is regulated during CP, but not during adult enhanced plasticity. Among downregulated DTGs, nuclear localization, heterochromatin, and DNA repair pathways are prominent. Accordingly, upstream TF activity cluster C2 supports chromatin reorganization and p53 signaling at P30, as several of its components modulate the p53 axis^30,40,41^. DNA repair modules include MRN complex-related signaling, BER-associated components, repair complex chaperones, and DSB sensing proteins, all of which are classically linked to DSB formation. These DTGs are coherent with the phenotypic change in γH2Ax foci observed in adult PV cells.

Interestingly, the higher number of γH2Ax foci at P30 is similar to the increased foci number observed after oxidative stress in dopaminergic neurons, which resulted in the disruption of a large intense γH2Ax foci into smaller ones^42^. Several studies have proposed that sustained neuronal activity and/or IEG-associated transcription can be coupled to γH2Ax-marked DSBs, a process that could happen without external stimulus during the CP^43,44^. An unresolved point is whether heightened CP plasticity contributes to DSB formation or, conversely, if DSB formation reflects a chromatin state prone to developmental plasticity. Our findings of DTGs associated with decreased DNA repair response suggest that CP plasticity could represent a period prone to DSBs. The question of whether increased number of DSBs is harmful or is physiologically tolerated remains an open question, as DSBs are associated with neurodegeneration^45^, but are not necessarily a driver of it^46^. Finally, the higher number of H3K9me3 and DAPI foci at P30, along with DTGs related to DNA and histone modifications, suggest that the CP initiates heterochromatin restructuring in PV cells. These observations are coherent with the gradual decline of MeCP2 foci in WFA^+^ cells after CP in either V1 or the auditory cortex^31^.

Interestingly, adult plasticity via ChABC treatment does not recapitulate the chromatin-linked phenotype. While subtle or transient nuclear effects cannot be excluded, this separation supports cautious optimism regarding translational safety, as enhancing plasticity in the adult brain has become a broadly pursued therapeutic. A single injection of ChABC, as performed in the present study, is sufficient to enhance OD plasticity and restore motor function^22,47,48^, but either long-term infusion or viral delivery^49^ might be required in complex cases for accurate remodeling, in which case long-term impact on the PV cell translatome might differ. More broadly, plasticity-enhancing methods such as dark exposure, environmental enrichment, pharmacological manipulations, or growth factor-based strategies, are increasingly considered in translational contexts^6,7,22,48,50,51^, and would benefit for more precise characterization of their effects on the PV translatome.

### Study limitations

The choice to analyze ribosome-associated transcripts leaves open the question of the rate mRNA translation and protein turnover. This could be addressed essentially by a proteomic approach which is beyond the scope of the study. The difference in TRAP response between ChABC treatment and scFv-OTX2 expression is curious given that a decrease in OTX2 import downregulates PNN assembly and, reciprocally, that ChABC treatment reduces OTX2 internalization^52^. We predicted to see more DTGs in common between the two adult plasticity paradigms. The choice of one timepoint in the juvenile and adult mice may lead to some simplification of the phenomena. To analyze these paradigms with more granularity, DTG analysis could be extended beyond P30 (compared to P90), and at different times after ChABC injections and scFv-OTX2 expression. Similarly, the spatial identity of analyzed cells might also be important. In this study, we analyzed PV cell translatome across all cortical layers within V1. Although the fast-spiking basket cell represents most PV^+^ interneurons in V1 and the supragranular layer PV cells participating in CP regulation are well represented^53^, our analysis may be diluted by ribosome-associated mRNA from PV cells not involved in V1 plasticity mechanisms.

## Acknowledgements

We would like to thank Jessica Apulei and Uma Mani for image analysis. Funding was provided by the Neuroglia Fund and the Agence Nationale de la Recherche (ANR-22-CE16-0033-01) for supporting research costs and salaries. CC was granted a fellowship by the Major Research Program "PSL-NEURO", launched by PSL Research University and implemented by ANR (ANR-10-IDEX-0001).

## Methods

### Mice and surgery

The C57Bl/6J mice were purchased from Janvier Laboratories, while *PV:Cre* mice (strain #008069) and *NuTRAP* mice (strain #029899) were obtained from Jackson Labs, USA. The PV-NuTRAP mice were generated by crossing *PV-Cre* females and *NuTRAP* males to avoid spurious recombination in the testis, which have been shown to transiently express PV^54^. Recombinant ChABC (Bio-Techne) was injected in V1 (lambda: x = ±1.7 mm, y = 0 mm, z = 0.5 mm) with a single dose of 0.5 μl (50 U/mL) at a rate of 0.1 μl/min using a Hamilton 1701 syringe mounted with a 33G needle. All animal procedures, including housing, were carried out in accordance with the recommendations of the European Economic Community (2010/63/UE) and the French National Committee (2013/118). For surgical procedures, animals were anesthetized with xylazine (Rompun 2%, 5 mg/kg) and ketamine (Imalgene 500, 80 mg/kg). For histological and biochemical analysis, mice either underwent transcardial perfusion or were sacrificed by cervical elongation.

### Immunohistochemistry

Mice were perfused transcardially with PBS followed by 4% paraformaldehyde prepared in PBS. Brains were postfixed for 24 h at 4 °C and immunohistochemistry was performed on cryosections (20 μm) encompassing the entire visual cortex. Heat-induced antigen retrieval in 10 mM sodium citrate was performed prior to overnight primary antibody incubation at 4 °C. Primary antibodies and lectins included anti-PV (rabbit, Swant 27), anti-GFP (rabbit, abcam ab290), anti-γH2AX (mouse, 1/200, Millipore clone JBW301), anti-H3K9me3 (rabbit, 1/200, abcam ab8898), and biotinylated WFA (1/100, Sigma L1516). Secondary antibodies were Alexa Fluor-conjugated (Molecular Probes).

### Image Analysis

Images were acquired with a confocal microscope (Zeiss 800) with ZEN software (Zeiss). Image acquisition was performed at ×40 magnification, and a custom Java macro was developed for downstream image analysis. Cell bodies were segmented using a customized Cellpose deep-learning model trained on a dataset comprising 60 annotated examples (40 for training and 20 for testing), achieving an Intersection over Union (IoU) of 0.93, a precision of 0.982 and an accuracy of 0.971. This macro identified PV^+^ interneurons, WFA^+^ cells and GFP^+^ cells. Image analysis was implemented with a plugin for Fiji^55^, using the Bio-Formats^56^, CLIJ^57^ and 3D Image Suite^58^ libraries. The DAPI channel was downscaled 4-fold prior to nucleus detection with the 2D-stitched version of Cellpose^59^. Segmentation outputs were then rescaled to their original size, and 3D nuclei were filtered by volume to minimize false positives. PV^+^ and WFA^+^ cells were segmented similarly, using custom-trained Cellpose models (via the human-in-the-loop procedure). PV and WFA models were initialized with the ‘cyto’ and ‘livecell’ pretrained models, respectively, and fine-tuned on 40 manually-annotated images. Cells were assigned the nucleus associated with at least half of their volume. Cells without any associated nucleus were removed from the analysis. Each nucleus was thus labelled as positive or negative for PV and WFA, and the integrated intensity of each cell and associated nucleus was measured. To account for background noise, images were normalized by subtracting the median intensity value of the minimum intensity Z-projection for each channel. yH2AX, DAPI and H3K9me3 foci were detected in nuclei using StarDist 2D^60^, with a model trained on a dataset of 1950 labelled puncta. The integrated intensity of yH2AX and DAPI foci was measured. Channels were downscaled 2-fold before detection, then rescaled, labelled in 3D, and volume-filtered. The accuracy of the plugin was validated on a subset of 20 representative images. All scripts are available upon request.

### Optical imaging

For monocular deprivation, mice were anesthetized with a mix of ketamine (95 mg/kg) and xylazine (12 mg/kg) in 0.9 % NaCl (i.p.). The area surrounding the contralateral eye (left eye) was trimmed and wiped with 70% ethanol prior to performing two mattress sutures. Animals were checked daily to ensure sutures remained intact. After 3 days, mice were anesthetized with urethane (1.2 g/kg, i.p.), sedated with chlorprothixene (8 mg/kg, i.m.), and treated with Atropine (0.1 mg/kg, s.c.) and dexamethasone (2 mg/kg, s.c.). Optical imaging was performed as previously described^12^.

### Animal behavior

For the visual water maze, the apparatus consists of a pool (1.5 m x 80 cm) with 2 monitors placed side by side at one end. A 41cm midline divider is extended from the wide end into the pool, creating a maze with a stem and 2 arms. An escape platform invisible to the animals is placed below the monitor, where the grating is projected. The position of the grating and the platform is alternated in a pseudorandom sequence over the training and test trials. For measuring visual acuity, animals are initially trained to distinguish a low spatial frequency grating from a gray screen with the same brightness. Once 70% cumulative accuracy is achieved, the testing phase determined a visual acuity threshold by increasing the spatial frequency of the grating. During the first ten trials, grating frequency was increased after every successful trial and lowered after two missed trials. During the second phase, from ten trials to plateau, difficulty was increased whereby grating frequency was increased after two consecutive successes and decreased after one missed trial. Mice were given 10 trials a day, with 5 min in between intervals of rest, under a 50 W heating lamp.

The visual cliff test was performed using a transparent acrylic chamber (50 × 50 cm) positioned such that half of the apparatus rested on an opaque support platform, while the remaining half extended beyond the edge, creating the visual illusion of a drop. Mice were initially placed on the supported (solid) side of the chamber and allowed to freely explore the apparatus. Behavior was recorded using a fixed overhead camera. Video data were analyzed offline by an experimenter blinded to experimental conditions using ezTrack software. The total time spent in the apparent “void” area versus the solid area was quantified.

### TRAP protocol

Brain V1 cortices (bilateral) of 3-5 PV-NuTRAP mice per sample were manually dissected with minimal white matter and placed into a pre-chilled 2 ml dounce containing 1 ml cold lysis buffer (50 mM TrisHCl, pH 7.5, 100 mM KCl, 12 mM MgCl_2_, 1% NP40, 1mM DTT, 100 µg/ml cycloheximide (Sigma), 0.4 U/µl RNAse OUT, 1X Protease inhibitor (Pierce)). Homogenization involved 10 strokes with a loose pestle, 10 strokes with a tight pestle, 5 min incubation on ice, and 10 more strokes with a tight pestle. Homogenates were filtered on 40 μm cell strainer and transferred to a pre-chilled 2 ml LoBind tube. Homogenates were subsequently centrifuged at 4 °C, 10 000 rcf for 10 min. A 50 μl sample of supernatant was kept as input, and the remaining supernatant was transferred to a pre-chilled 1.5 ml LoBind tube. DynaBeads Protein G (Thermofisher, 1003D) were blocked in 1 mL lysis buffer supplemented with 2% BSA and 0.2 mg/ml yeast tRNA for 3 x 1 min incubation, and resuspended in their original volume. A 10 min pre-clear step was performed with 10 µl DynaBeads Protein G per sample at 4°C. Beads were removed with a Dynamag magnet (Thermofisher) and samples were incubated with 1 µg anti-GFP antibody (ab290, abcam) for 1h at 4°C. 30µL Dynabeads Protein G were added to each sample and incubated 2h at 4°C. Samples were placed on a DynaMag magnet for 1 min until complete clearing of the beads in solution. Beads were washed 3 times with high salt lysis buffer (50 mM TrisHCl, pH 7.5, 300 mM KCl, 12 mM MgCl_2_, 1% NP40, 2 mM DTT, 100 µg/ml cycloheximide, 0.4 U/µl RNAse OUT, 1X Protease inhibitor). After the third wash, samples were removed from the magnet and beads were resuspended in 500 µL QIAzol and kept at -80 °C until RNA extraction using RNeazy Micro Kit (Qiagen) with standard procedure. Double DNase (Zymo Research) treatment was performed. Purified RNA was quantified on a Qubit 4 and analyzed on a BioAnalyzer for quality control.

### Sequencing & Bioinformatic analyses

Samples were sequenced by the Curie Institute Genomics facility with the SMARTer Stranded Total RNA-Seq Kit (Pico Input Mammalian). Bioinformatic analysis was conducted using a custom pipeline. Articles and packages are provided in the tools section. Custom scripts are available upon reasonable request.

All computational analyses were performed in R (v4.3.3). Raw gene-level count matrices and sample metadata were imported and harmonized with gene annotations. Samples were encoded with an explicit factor combining fraction (PV-IP or Input) and experimental condition and were interrogated through three contrast families: PV-IP versus Input within each condition, PV-IP versus PV-IP and Input versus Input.

Differential expression testing was conducted with the edgeR quasi-likelihood negative binomial GLM framework^61^. Library sizes were normalized using calcNormFactors with the TMMwsp method (edgeR implementation of TMM-style scaling). Lowly expressed genes were filtered using min.count = 30, min.total.count = 100, and min.prop = 0.6 in the primary model fits. Highly variable mitochondrial (mt-), immunoglobulin (Ig) or major histocompatibility (H2-) prefixes were removed. Dispersion estimation and model fitting used robust procedures (estimateDisp(robust = TRUE), glmQLFit(robust = TRUE)), and contrasts were tested using quasi-likelihood F-tests (glmQLFTest). Statistical significance was defined as FDR < 0.05 using the Benjamini–Hochberg procedure.

Pulldown quality was assessed in the PV-IP vs Input comparison, with the enrichment of *Pvalb* versus other cells markers such as *Gfap* and *Cxc3cr1*. The first threshold of low enrichment was set at mean_log2FC > -1.36 to filter out glial cells markers that likely correspond to background (∼10% of detected genes, 1200 genes). Then, within each condition, a second threshold at log2FC > 0.3 with FDR < 0.05 was set to define PV-enriched transcripts.

Set-overlap summaries and visualizations were computed using RVenn and UpSetR^62^. Across conditions, we constructed three sets: a core (intersection of all condition-specific PV-enriched sets), condition-specific sets (genes enriched in each condition but in none of the others), and a union (genes enriched in at least one condition).

After removal of lowly enriched genes, all PV-IP fractions were compared to the P90 PV-IP fraction that represent the stable adult baseline. In all differential-expression gene lists, the primary effect-size threshold was |log2FC| ≥ 0.5 with FDR < 0.05, corresponding to the PV-IP Filtered fraction.

### Transcription factor activity inference

Upstream regulator analysis identifies TF based on the confidence of interaction between TF-gene pairs (regulons) and level and directionality of differentially expressed genes. The mouse DoRothEA resource was used to retain high-confidence regulons (confidence A/B/C, removing D/E)^63^. For each TF, VIPER was run on the voom-normalized PV-enriched expression matrix to obtain a sample-by-sample TF activity matrix^64–66^. Hierarchical clustering was performed on the predicted TF activity across all conditions^67^. To address regulatory consistency for each TF, “leading-edge” target genes were identified by correlating each target’s expression across PV samples with the TF’s activity and combining with the target’s mode of regulation ∈ {−1,+1}; targets with positive signed evidence (mor × correlation > 0) were prioritized. For each TF, we reported and visualized only leading-edge targets, annotating each target with its expected direction (up or down) given the TF’s activity and mode of regulation.

### Gene ontology over-representation analysis

For background universe construction in enrichment analyses, expressed-gene universes were defined for PV-IP as genes with counts > 30 in at least two samples (nMinExpr = 30, nSamples = 2). Gene Ontology enrichment analyses were performed using the topGO package with the weight01 algorithm and Fisher’s exact test, which accounts for the hierarchical structure of the GO graph and down-weights broad parent terms. Gene sets were defined from the PV-filtered differential expression tables, to avoid splitting pathways (i.e., a pathway comprised of PV-enriched genes and housekeeping genes below the enrichment threshold). Genes were separated as ALL, UP, and DOWN subsets for each contrast, together with ‘common’ and contrast-specific masks (Common corresponding to the concordant intersection of PV_P30 and PV_ChABC, specific sets to their respective unique genes). Splitting Up and Down provides a better biological interpretability, at the expense of p-value strength^68^. The background universe was restricted to the PV-expressed genes (i.e genes expressed at least 30 counts in half of the PV samples), ensuring comparable annotation depth. For each ontology (Biological Process, Molecular Function, Cellular Component), topGO data objects were built using the org.Mm.eg.db annotation (https://doi.org/doi:10.18129/B9.bioc.org.Mm.eg.db) with Ensembl identifiers. Terms were retained when annotated to between 6 and 500 genes to avoid very large unspecific pathways (such as “Membrane”) or pathways driven by one single gene. The output included GO identifier, description, ontology, p-values, term size, number of significant genes, and the list of gene symbols.

### Statistical Analysis

Statistical analysis of immunohistochemistry, optical imaging, and animal behavior were performed with either pairwise comparisons using Student t-test or multiple group analyses using ANOVA (one- or two-way) followed by Bonferroni correction (Prism 9, GraphPad). Changes in frequency distribution were assessed by Kolmogorov–Smirnov test. Specific tests are indicated in Figure legends.

---

**Supplementary Fig. 1.**
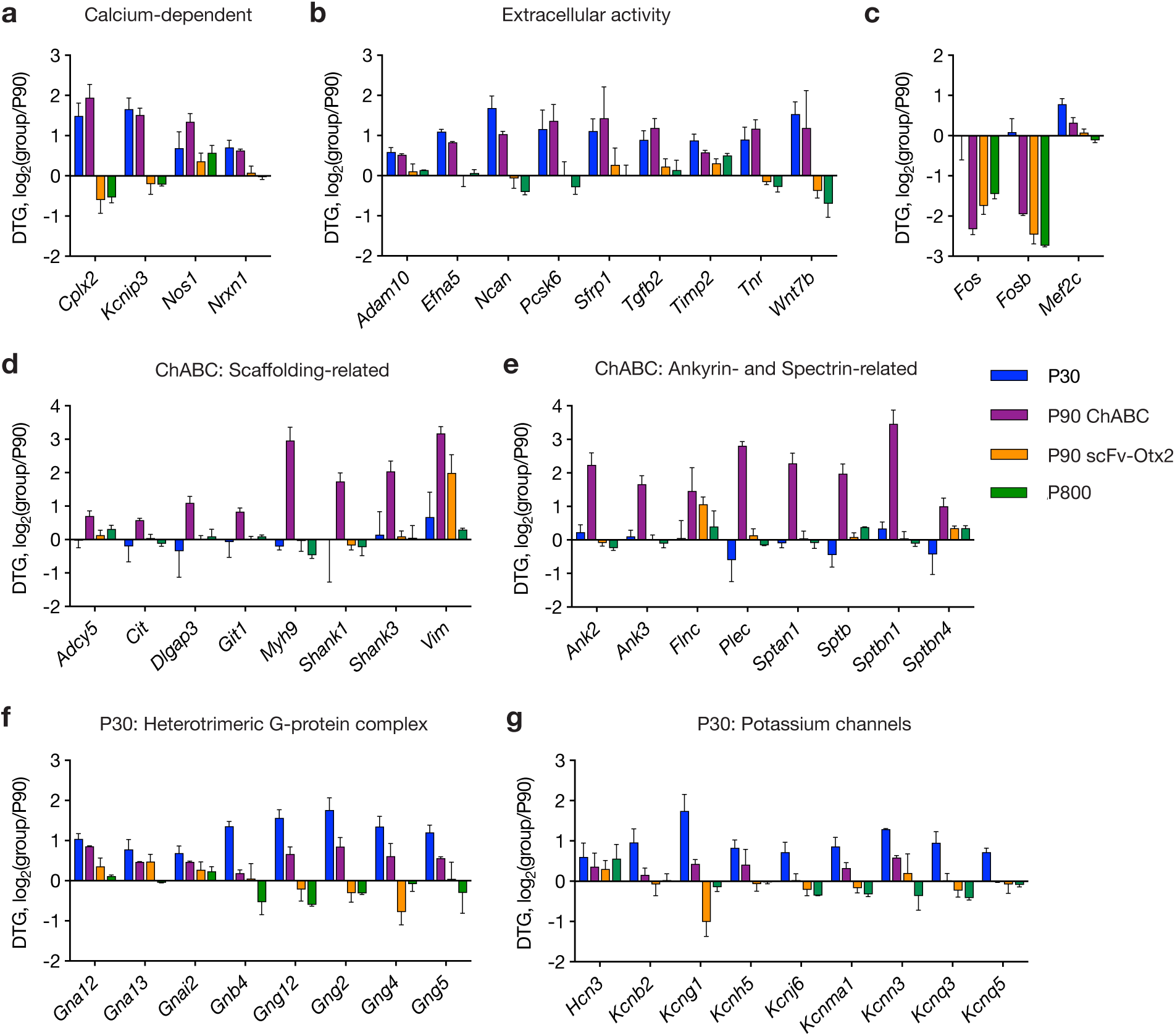
Comparison of DTGs selected for genes of interest in the P30 and P90 ChABC groups. **a-b,** Sample of DTGs with similar responses in both P30 and P90 ChABC groups for proteins with either calcium-dependent activity (**a**) or extracellular activity (**b**). **c,** Sample of DTGs for transcription factors and IEGs of interest. **d-e,** Sample of DTGs with specific responses in the P90 ChABC group for proteins related to scaffolding (**d**) or ankyrin and spectrin functions (**e**). **f-g,** Sample of DTGs with specific responses in the P30 group for heterotrimeric G-protein complexes (**f**) or potassium channels (**g**).

